# Protein Fold Classification using Graph Neural Network and Protein Topology Graph

**DOI:** 10.1101/2022.08.10.503436

**Authors:** Suri Dipannita Sayeed, Jan Niclas Wolf, Ina Koch, Guang Song

## Abstract

Protein fold classification reveals key structural information about proteins that is essential for understanding their function. While numerous approaches exist in the literature that classifies protein fold from sequence data using machine learning, there is hardly any approach that classifies protein fold from the secondary or tertiary structure data using deep learning. This work proposes a novel protein fold classification technique based on graph neural network and protein topology graphs. Protein topology graphs are constructed according to definitions in the Protein Topology Graph Library from protein secondary structure level data and their contacts. To the best of our knowledge, this is the first approach that applies graph neural network for protein fold classification. We analyze the SCOPe 2.07 data set, a manually and computationally curated database that classifies known protein structures into their taxonomic hierarchy and provides predefined labels for a certain number of entries from the Protein Data Bank. We also analyze the latest version of the CATH data set. Experimental results show that the classification accuracy is at around 82% − 100% under certain settings. Due to the rapid growth of structural data, automating the structure classification process with high accuracy using structural data is much needed in the field. This work introduces a new paradigm of protein fold classification that meets this need. The implementation of the model for protein fold classification and the datasets are available here https://github.com/SuriDipannitaSayeed/ProteinFoldClassification.git

**Author summary:** Classification of protein structures is traditionally done using manual curation, evolutionary relationship, or sequence comparison-based methods. Applying machine learning and deep learning to protein structure classification is a comparatively new trend that holds great promises for automating the structure classification process. Advance deep learning technique like Graph Neural Network is still unexplored in this respect. SCOP and CATH are two traditional databases that provide the hierarchical taxonomic classification of protein structures. This work provides a novel computational approach that classifies protein folds in SCOP and CATH with graph neural network, performing a graph classification task.

## 1 Introduction

A protein fold is a 3D pattern in proteins that share the same composition in secondary structure elements as well as the same spatial arrangement and topology. Protein fold recognition is a problem of predicting protein folds from a given amino acid sequence or from a given protein structure. The problem of protein fold recognition is highly important for multiple reasons. Recognition of protein fold could be the first step to an efficient prediction of protein structures from amino acid sequences. On the other hand, protein structure classification is an important task also in structural biology, where newly determined structures are constantly needed to be assigned to their corresponding folds. Given the growing number of high resolution structure data in the Protein Data Bank (PDB) ^1^ and other data sources ^2^, a fast, accurate, and fully automated approach for structure classification procedure is much needed.

Traditionally, sequence-based protein fold recognition methods can be divided into two categories. Those are: 1) sequence alignment-based methods ^3^ and 2) machine learning-based *ab-initio* methods ^4^. In sequence alignment-based methods, a query protein sequence or sequence profile is aligned against templates with similar structures to identify the most probable fold. Machine learning models from the very beginning time such as support vector machines ^4^, kernel-based methods and ensemble classifiers ^5^ also rely on sequence correlation-based features. These models work fine in recognizing folds at the family level, where sequences have enough evolutionary relationships. But the problem of fold recognition becomes significantly more challenging when sequences are highly dissimilar and there is not enough evidence of evolutionary relationships among sequences. This indicates the necessity to integrate structural features in the prediction.

The Structural Classification of Proteins (SCOP) ^6^ and Hierarchical Classification of Protein Domain Structures (CATH) ^7^ are two standard databases for taxonomic classification of protein domain structures. In both of these databases, protein structures are classified manually for the most part by considering their evolutionary relationships with other similar structures. SCOP classification is done mainly based on evolutionary relationship between homologous sequences. SCOPe ^8^ provides an automated pipeline for structure classification at protein family-fold level and superfamily-fold level. The basis of their classification approach is that, every protein has structural similarities with some other proteins that share a common evolutionary origin. Protein structures are grouped into homologous families based on their sequence similarity. SCOPe classification starts with the domain. The domains are then classified into the following hierarchy:

- Species: represents a unique protein sequence and its variants.
- Protein: groups together similar sequences with the same functions that either come from different biological species or from different isoforms within the same species.
- Family: combines proteins with similar sequences but distinct or similar functions. Due to their sequence similarity and or functional similarity, protein domains that are clustered in same family are likely to have a common evolutionary origin.
- Superfamily: merges together protein families with similar functions and structural features and come from a common evolutionary ancestor.
- Fold: is the purely structural layer of this classification. Structurally similar superfamilies are combined into folds.
- Class: combines folds based on their secondary structure content and their topological arrangements.

CATH ^7^, on the other hand, considers structural similarity over evolutionary relationships in classifying protein structures. Evolution gives rise to families of structurally similar proteins for which there is no evident evolutionary relationship as they have extremely low sequence similarities. CATH applies a semi-automatic approach for this classification. They use the standard multiple sequence alignment method and the structure comparison algorithm SSAP (Sequential Structure Alignment Program) ^9^. CATH has four levels in their hierarchy of classification:

- Class, C-level: describes the secondary structure element composition of each domain. It is the first level of the hierarchy.
- Architecture, A-level: describes the overall shape of the structure. This shape is revealed by the composition and orientation of the secondary structure units. At this level the connectivity among the secondary structure elements are not considered.
- Topology (Fold family), T-level: At this level, structures are clustered into fold families based on their overall shape and connectivity of secondary structure elements. Some fold families are overpopulated than others, and CATH organizes those by the SSAP score.
- Homologous Superfamily, H-level: When structurally similar proteins coming from the same T-level also have similar functions, they are supposed to share a common evolutionary ancestor. These proteins are grouped into a homologous superfamily.

The construction process of CATH is more automatic than that one in SCOPe. Both of these two databases have been considered as gold standards for benchmarking automatic pipelines of structure classification and for training machine learning-based models.

Every protein structure is shaped by the arrangement of secondary structure elements. And each structure defines a particular conformation in the conformation space. Recent revolution of deep learning has opened up opportunities for exploring more structural information, such as secondary structure composition and inter-residue contacts, in fold classification. Early deep learning-based models like DN-Fold ^10^ that relied upon evolutionary distance between protein pairs showed promising performance in classifying protein folds at family and superfamily level but very limited performance at the fold level. Sequence-based classification techniques such as DeepSF ^11^ that use Deep Convolutional Neural Network (DCNN) can classify protein folds with an accuracy of around 75.3%. Fold recognition method like DeepFR ^12^ exploits inter-residue contact information using DCNN. Inter-residue contact information holds important information about the overall 3D structure of the protein. Residue contacts can be local or non-local. Local contact information holds information about the local structural motifs, whereas non-local interactions among residues give information about the overall arrangements of the motifs. Both of these information could be valuable for recognizing the overall protein fold. Classification techniques like PRO3DCNN ^13^ also use DCNN to exploit fold-specific features. In addition to using a regular contact distance matrix, they use also one extra feature that is called persistance homology image of the protein structure. In combination with these two features, they reached an accuracy of around 95% − 98% for their dataset.

In this work, we propose a novel graph neural network (GNN)-based model to classify protein folds. GNN ^14^ is effective in capturing both local interactions and global interactions. The intuition behind using GNN is that it captures information from both local structural motifs and non-local residual contacts to generate a better *embedding* for classification. We use the open-source implementation of HGP-SL (Hierarchical Graph Pooling with Structure Learning) ^15^ for performing the graph classification task. The model takes protein topology graphs as input. The input graphs are constructed according to the definitions of Protein Topology Graph Library (PTGL) ^16^. We consider three node features for our classification, namely, i) node degree, ii) node type, and iii) the number of residues in the node (each node represents a secondary structure element). Our model has two basic layers: i) graph convolution layer and ii) hierarchical graph pooling layer. We apply a sophisticated structure learning mechanism with the graph pooling layer. At the very end of the model there is a softmax layer that generates a graph embedding to classify the graph against multiple classes. In our case, the class labels are the fold labels. We will discuss our methods and results in Sections *Material and Methods* and *Results and Discussion*, respectively.

## 2 Contribution of This Method

We claim that for protein structure classification, a highly accurate machine learning-based model that learns pattern from structural data provides two major benefits. We discuss them in details below.

### 2.1 Automating the Structure Classification Process

We focus particularly on classifying the structural layer that is the fold of the protein structure taxonomy. Due to the abundance of structure data, it seems that time is ripen for fully automating the process using deep learning.

- **Abundance of structure data solved at high-resolution**. Due to the advancement of technology in both hardware and software, the number of protein structures solved at high-resolution is increasing rapidly. Over the past 60 years X-ray crystallography has been the primary method for structure determination. A major limitation of X-ray crystallography is that it requires molecules to be crystallized, which is not always possible. Figure 1 shows the overall statistics of the growth in the number of structures in the Protein Data Bank (PDB). The data in PDB are generally collected from X-ray crystallography, NMR spectroscopy, or cryo-electron microscopy. Nuclear magnetic resonance (NMR) technique determines the 3-D structures of macromolecules in solution. This technique is limited mostly to small to medium macromolecules.

**Figure 1:**
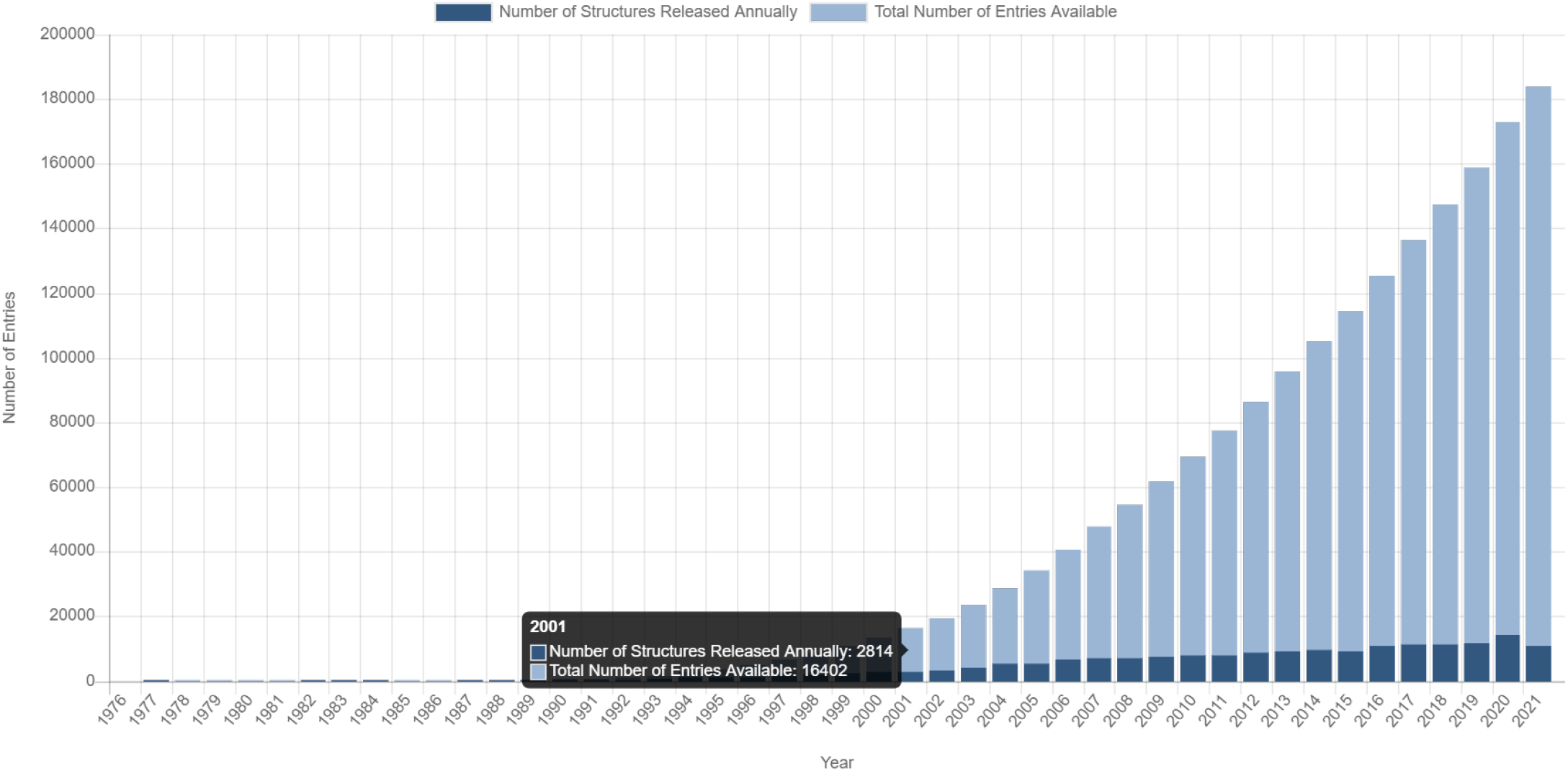
PDB Statistics: overall growth of released structures per year. https://www.rcsb.org/stats/growth/growth-released-structures
- Apart from PDB, Electron Microscopy Data Bank (EMDB) has also become a great resource for highresolution protein structure data. Currently, cryo-EM is one of the most prominent approaches for determining high-resolution structures of small to large macromolecular complexes (Figure 2).

**Figure 2:**
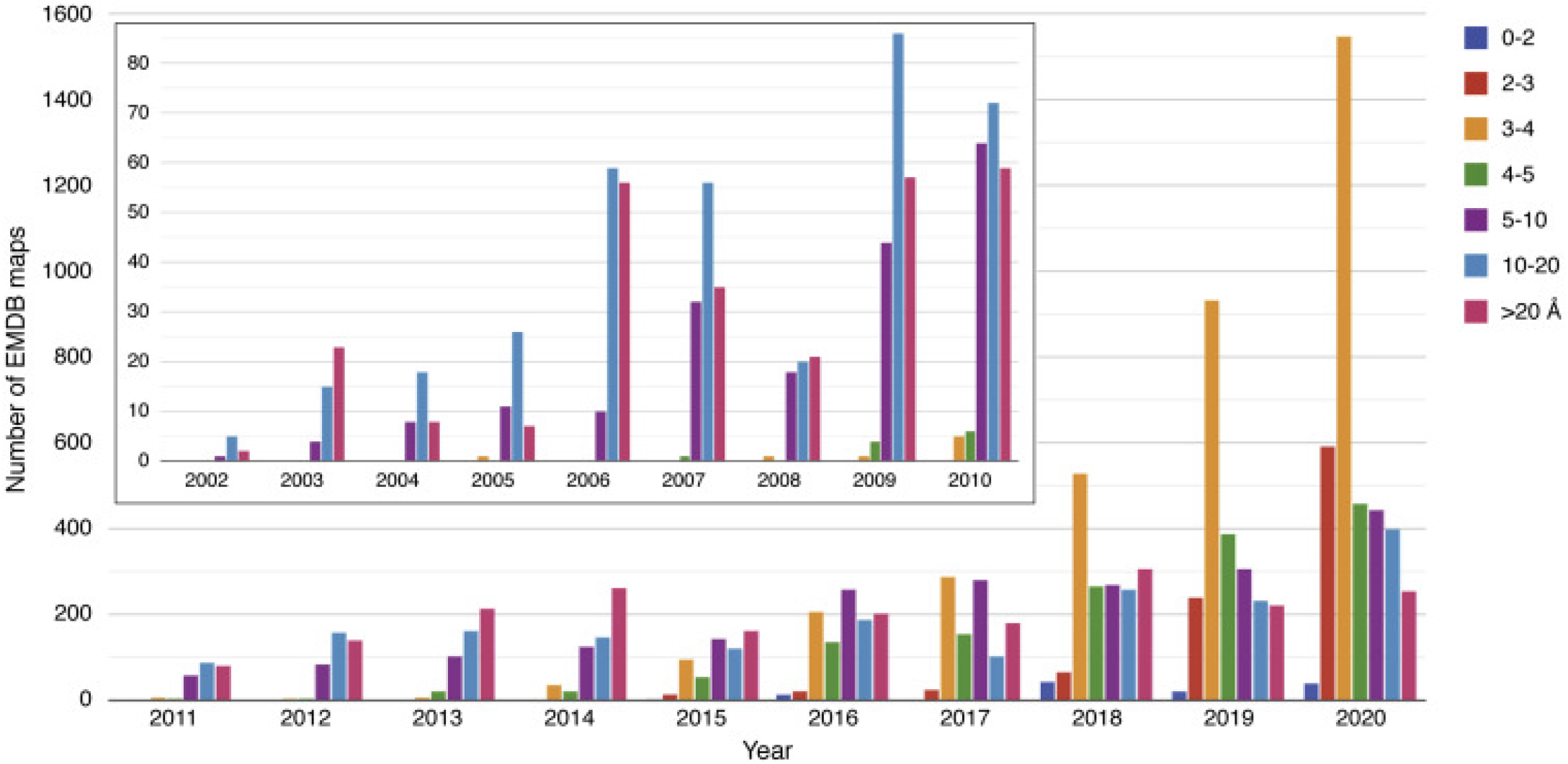
EMDB Statistics: overall growth of archived electron microscopy density maps at different resolution ranges between 2002 and 2020. https://ars.els-cdn.com/content/image/1-s2.0-S0021925821003380-gr6.jpg
- Apart from these experimental approaches, artificial intelligence(AI)-based systems such as AlphaFold ^17^ is also making a big leap towards generating highly accurate protein structure data at high throughput. AlphaFold has generated accurate protein structure predictions since its inception and has now achieved an accuracy that is comparable to experimental methods. AlphaFold database ^18^ is a fast growing resource of predicted protein structures. Currently, it contains protein structures from human proteome and 20 other key organisms. These structures are available to the scientific community and can be accessed through the UniProt accession ID. One important aspect about the newly predicted structures is that they are unclassified. With the ever increasing computationally-predicted structures, a deep learning-based automated method that can accurately classify them into their respective folds seems most consistent and natural.

### 2.2 Classifying Non-homologous Proteins

Sequence similarity-based fold classification techniques perform poorly when sequence similarity is low. Due to events like gene duplication and gene speciation, many sequences diverge from their homologs but their structures and functions remain mostly intact. Sequence-based models rely mainly on sequence homologs. There are many groups of proteins whose sequence similarity is lower than the required threshold but share significant structural similarity. Such structural similarities in the absence of any evident evolutionary relationship can be explained only by the similarities among the packing arrangement of the secondary structure elements. Folds like the globin-like family from the all-alpha architecture contain protein pairs whose sequence identity is less than 15% but share similar structures and functions. Figure 3 shows the similarities between two globin-like structures, and Figure 4 displays their similarities in terms of topology graph ^19^. The two graphs share a high degree of similarity in terms of the number of secondary structure elements and their contacts.

**Figure 3:**
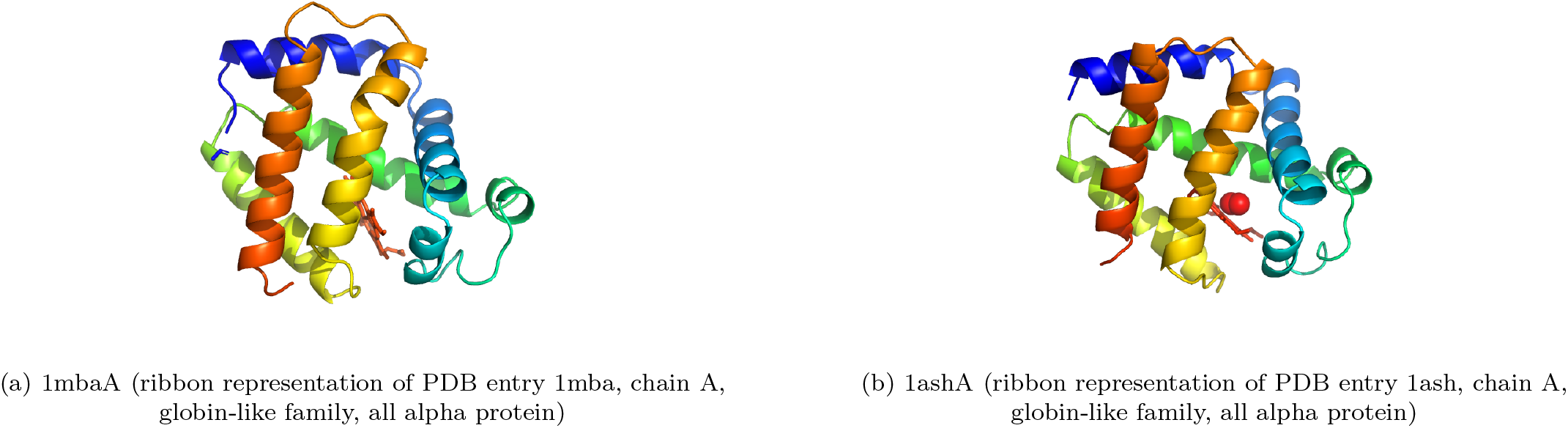
PyMOL structures of a similar-fold protein pair with sequence identity less than 15%

**Figure 4:**
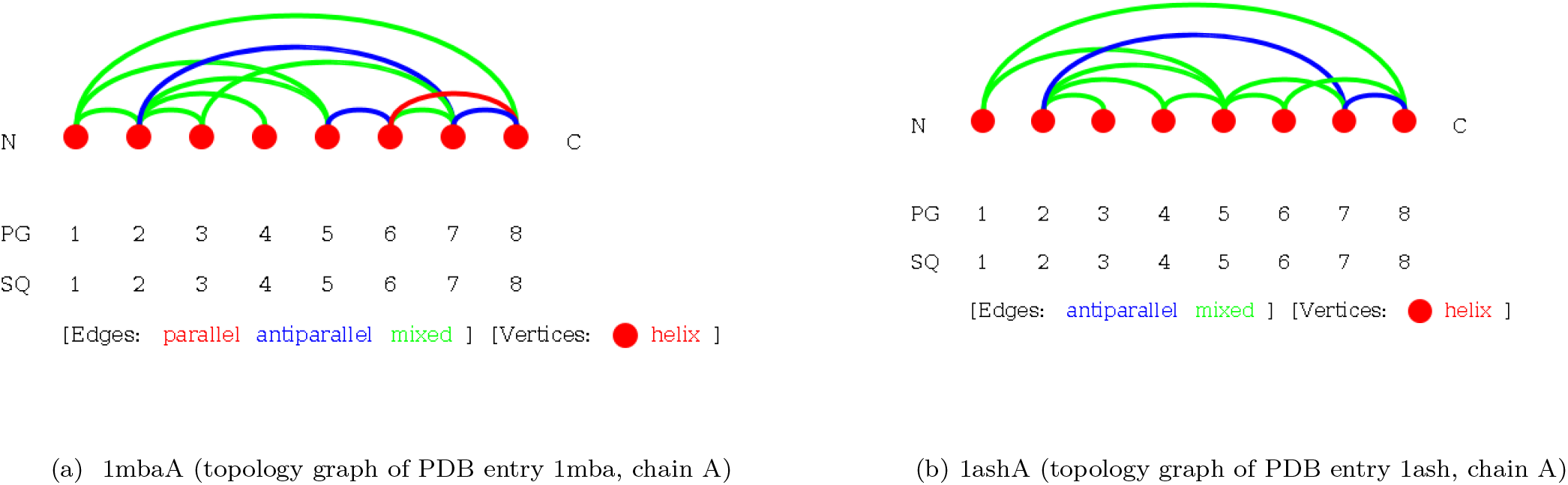
Topology graphs of a similar-fold protein pair with sequence identity less than 15%

The Tim-barrel fold/topology from the alpha-beta class, barrel architecture also displays a high degree of structural similarity in the absence of any evident sequence similarity. This fold is also challenging to classify using sequence similarity-based machine learning models. Figure 5 shows the PyMOL structures of two examples from the Tim-barrel fold. Their topology graphs are given in Figure 6.

**Figure 5:**
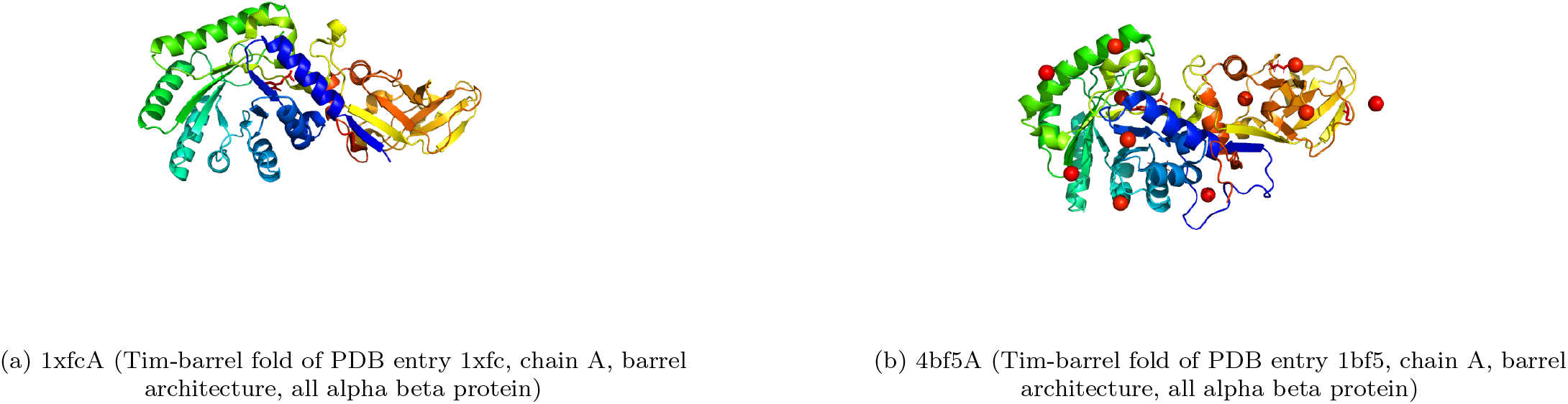
PyMOL structures of two similar-fold proteins with a sequence identity less than 30%

**Figure 6:**
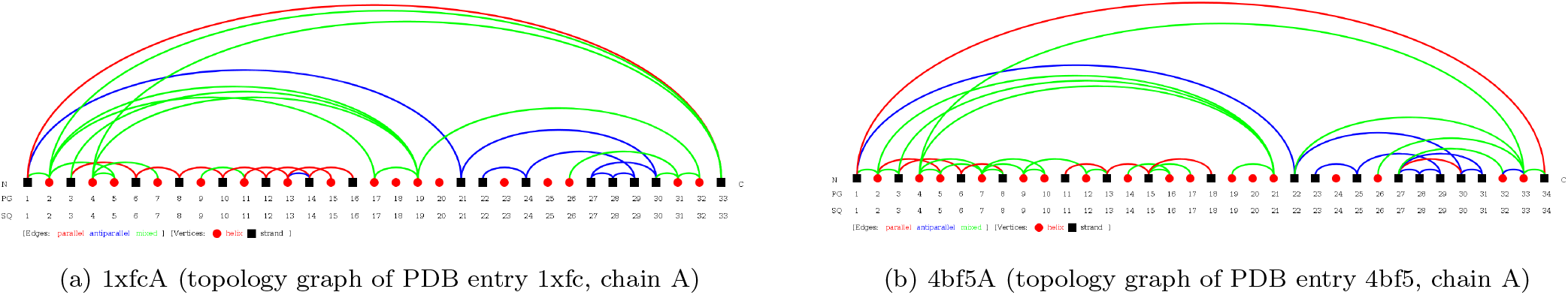
Topology graphs of two similar fold proteins with a sequence identity less than 30%

The above examples show that a simple and intuitive approach based on protein topology graphs can potentially solve the problem with non-homologous protein pairs for protein fold recognition.

### 2.3 Classifying Against a Large Number of Classes

Classifying protein fold using only secondary structure elements and their topological connections is a more abstract but robust way of classification. In the cases of sequence-based approaches, when more classes are included in the data set, it becomes increasingly challenging to classify fold patterns accurately for a query protein sequence. Classifying against a large number of classes is challenging also for traditional classification models like Support Vector Machine (SVM) and Deep Neural Network (DNN). Our GNN-based approach can easily classify against multiple classes. For example, under Orthogonal Bundle architecture in mainly alpha class of the CATH dataset ^7^, there are 290 topologies. Our model can classify against these 290 classes as well as it does against a smaller number of classes.

In summary, these points underline the potential of our proposed method for automatic structural classification.

## 3 Results and Discussion

To corroborate our proposed method we first analyze small data sets like DD, EDD and TG ^20^. We find our graph-based learning model outperforms existing sequence based learning models on these benchmark data sets regarding the protein fold classification task. To further expand our analysis on small data sets, we train and test our model also on large-scale protein structure classification data sets: SCOPe ^8^ and CATH ^7^.

### 3.1 Results on Benchmark Data Set

In this section, a combination of *ab-initio* machine learning models are compared on three benchmark data sets; DD, EDD and TG ^20^. These models learn patterns mainly based on sequence profile-based features and some structural and chemical features. As our approach is graph learning-based, graphs are collected for the respective protein entries in these three data sets. After that it is trained and tested on the converted graph data sets. Ten-fold cross validation is applied on DD, EDD and TG data sets. The DD data set is also evaluated using the independent test set that comes along with the original data set. The classification accuracy is reported in Table 2.

**Table 1:** Comparison of results from different methods on three benchmark data sets

| Method | Accuracy |  |  |
| --- | --- | --- | --- |
|  | DD | EDD | TG |
| ACCFold<sup>21</sup> | 0.701 | 0.859 | 0.664 |
| TAXFOLD<sup>22</sup> | 0.715 | 0.9 | NA |
| PFPA<sup>5</sup> | 0.736 | 0.926 | NA |
| HMMFold<sup>23</sup> | 0.758 | 0.938 | 0.86 |
| NiRecor<sup>24</sup> | 0.812 | 0.917 | 0.846 |
| SVM-fold<sup>4</sup> | 0.773 | 0.945 | 0.865 |
| TA-fold<sup>4</sup> | 0.799 | 0.971 | 0.927 |
| ProFold<sup>25</sup> | 0.725 | 0.936 | 0.94 |
| DeepFrag-k<sup>26</sup> | 0.825 | 0.965 | 0.96 |
| HGP_SL (This Article) | 0.98<sup>a</sup> | 0.96<sup>a</sup> | 1.0<sup>a</sup> |
|  | 0.827<sup>b</sup> | — | — |

*a* from ten-fold cross validation accuracy, *b* from independent data set

**Table 2:** One-vs-all classification on DD data set

| # | Fold ID | Fold Name | DeepFrag-k Accuracy | ProFold Accuracy | HGP-SL Accuracy |
| --- | --- | --- | --- | --- | --- |
| 1 | a.1 | Globin-like | 98.0 | 100.0 | 100.0 |
| 2 | a.3 | Cytochrome c | 95.0 | 100.0 | 100.0 |
| 3 | a.4 | DNA?RNA-binding 3-helical bundle | 85.9 | 60.0 | 100.0 |
| 4 | a.24 | 4-Helical up-and-down bundle | 91.5 | 87.5 | 100.0 |
| 5 | a.26 | 4-Helical cytokines | 98.9 | 88.9 | 100.0 |
| 6 | a.39 | EF hand-like | 90.8 | 77.8 | 100.0 |
| 7 | b.1 | Immunoglobulin-like beta sandwich | 91.10 | 84.1 | 100.0 |
| 8 | b.6 | Cupredoxin-like | 78.7 | 66.7 | 100.0 |
| 9 | b.121 | Nucleioplasm-likeNP | 91.3 | 92.3 | 100.0 |
| 10 | b.29 | CpnA-like lectins/glucanases | 76.7 | 66.7 | 100.0 |
| 11 | b.34 | SH3-like barrel | 78.0 | 50.0 | 100.0 |
| 12 | b.40 | OB-Fold | 80.4 | 68.4 | 100.0 |
| 13 | b.42 | Beta-Trefoil | 89.0 | 100.0 | 100.0 |
| 14 | b.47 | Trypsin-like serine proteases | 75.0 | 50.0 | 100.0 |
| 15 | b.60 | Liocalins | 90.5 | 100.0 | 100.0 |
| 16 | c.1 | TIM beta/alpha barrel | 93.8 | 93.8 | 100.0 |
| 17 | c.2 | FAD/NAD(P)-binding domain | 89.7 | 91.7 | 100.0 |
| 18 | c.3 | Flavodoxin-like | 60.2 | 46.2 | 100.0 |
| 19 | c.23 | NAD(P)-binding Rossmann | 90.2 | 85.2 | 100.0 |
| 20 | c.37 | P-loop containing NTH | 79.5 | 50.0 | 100.0 |
| 21 | c.47 | Thioredoxin-fold | 97.5 | 87.5 | 100.0 |
| 22 | c.55 | Ribonuclease H-like motif | 75.3 | 58.3 | 100.0 |
| 23 | c.69 | alpha/bet-Hydrolases | 78.4 | 71.4 | 100.0 |
| 24 | c.93 | Periplasmic binding protein-like | 92.0 | 100.0 | 100.0 |
| 25 | d.15 | beta-Grasp(ubiquitin-like) | 69.4 | 25.0 | 100.0 |
| 26 | d.58 | Ferredoxin-like | 76.8 | 59.3 | 100.0 |
| 27 | g.3 | Knottins(small inhibitors, toxins, lectins) | 88.2 | 96.3 | 100.0 |

The DD data set was analyzed using one-vs-all classification scheme to compare the fold prediction accuracy of each individual class. Results are compared with two other methods; DeepFrag-K and ProFold. The results of DeepFrag-k and ProFold are collected from Ref. 26. Our method secures a uniformly high classification accuracy across different folds. There is a huge performance gap in classification accuracy for 18 out of 27 fold classes for other methods. For example, for folds such as SH3-like barrel, trypsin-like serine proteases, flavodoxin-like, P-loop containing NTH and ubiquitin-like, Profold demonstrates a low accuracy, whereas DeepFrag-K shows a significantly higher accuracy. HGP-SL shows 100% accuracy for all 27 folds. The evaluation of each class is also done using the independent test set available for the DD data set.

### 3.2 Results on SCOPe 2.07 Data Set

Protein Structures are grouped into four major classes in the SCOPe database ^8^, namely, alpha, beta, alpha/beta, and alpha+beta. These structural classes define the composition of secondary structure elements in SCOPe domains. The next level in the SCOPe structure classification hierarchy is fold. There are 289 folds in the all-alpha class, and 178, 148, and 388 folds in the all-beta, alpha/beta, and alpha+beta classes, respectively. In this work, we focus our classification problem at the fold level and aim to classify all protein domains under every structural class into their corresponding folds.

First, we collect topology graph data ^19^ from every structural class without applying any filtering. For the all-alpha data set, we have 49, 670 protein domain graphs. There are 86, 430, 96, 716, and 84, 473 graphs for all-beta, alpha*/*beta, and alpha+beta classes, respectively. These data sets reflect the diversity in the original PDB data source since every unique protein chain in every structural class is included. The data sets are trained for 5, 500 epochs using Adam optimizer and cross-entropy loss.

The ten-fold cross validation results on the data sets are shown in Table 3. From the results, it is seen that the average accuracy of fold classification by our model is around 90%. The accuracy of fold classification for the all-alpha class is slightly lower, at around 88%.

**Table 3:** Prediction accuracies across different structural classes of SCOPe domains using the unfiltered data set

| Class | No of fold groups | No of graphs | Accuracy(%) |
| --- | --- | --- | --- |
| All-alpha proteins | 289 | 49,670 | 88% |
| All-beta proteins | 178 | 86,430 | 93% |
| Alpha/beta proteins | 148 | 96,716 | 91% |
| Alpha+beta proteins | 396 | 84,473 | 92% |

Many sequences in the PDB are highly similar, or even identical, to one another. Sequences that share high sequence identity usually have the same fold. To investigate how sequence similarity in the data set influences the performance of our method, we examine in the following the dependency of classification accuracy on sequence similarity. To this end, we cluster structures at different sequence similarity levels. We use the service provided by PDB sequence clusters data. The data, updated weekly, provides clusterings of all the protein chains in the PDB by MMseqs2^27^ at sequence identity levels of 30%, 40%, 50%, 70%, 90%, 95%, and 100%, respectively. In MMseqs2, protein sequences are clustered based on sequence similarity thresholds. Each protein chain is assigned to one and only one specific cluster, with sequences within the cluster share a sequence similarity equal to or greater than the threshold. As the threshold increases from 30% to 100%, the number of clusters increases. To keep the sequence similarity at a certain identity level, only one chain is randomly selected from each cluster for classification. The folds are classified under all four structural classes, and at different sequence identity levels. The model is trained for 3, 500 epochs with Adam optimizer and cross-entropy loss. The ten-fold cross validation results are given in Table 4. Note specifically that when the sequence identity threshold is at 100% (the bottom row of Table 4), the numbers of graphs are still greatly reduced when compared to those in Table 3 since identical sequences are filtered out from the data sets.

**Table 4:** Classification accuracy at different sequence identity levels for all-alpha, all-beta, alpha+beta, alpha/beta classes. #F and #G denote the number of folds and the number of protein domain graphs, respectively, while Acc denotes accuracy.

| Class | Alpha |  |  | Beta |  |  | Alpha+Beta |  |  | Alpha/Beta |  |  |
| --- | --- | --- | --- | --- | --- | --- | --- | --- | --- | --- | --- | --- |
| Seq. Sim. level | #F | #G | Acc | #F | #G | Acc | #F | #G | Acc | #F | #G | Acc |
| 30% | 289 | 3818 | 83% | 178 | 3964 | 98% | 396 | 4921 | 94% | 148 | 4582 | 98% |
| 40% | 289 | 4295 | 82% | 178 | 4840 | 97% | 396 | 5832 | 93% | 148 | 6238 | 96% |
| 50% | 289 | 4808 | 93% | 178 | 5531 | 92% | 396 | 6617 | 92% | 148 | 7709 | 96% |
| 70% | 289 | 5683 | 88% | 178 | 6814 | 97% | 396 | 7955 | 91% | 148 | 9788 | 95% |
| 90% | 289 | 6384 | 89% | 178 | 9081 | 97% | 396 | 9019 | 90% | 148 | 10973 | 94% |
| 95% | 289 | 6663 | 92% | 178 | 10463 | 97% | 396 | 9414 | 90% | 148 | 11337 | 94% |
| 100% | 289 | 10044 | 91% | 178 | 17145 | 96% | 396 | 14831 | 88% | 148 | 18329 | 94% |

In constructing topology graphs of protein domains, domain boundaries are considered. Whether a protein chain is multi-domain or single-domain, folds are classified at the domain level.

Table 4 presents the results for SCOPe 2.07 dataset at different similarity levels. Remarkably, while as the sequence similarity level is reduced from 100% to 30% and the number of graphs decreases, the classification accuracy increases and reaches even 98% for Beta and Alpha/Beta classes. This implies the removal of redundant sequences actually helps training and improves the performance. For one, the removal of redundant sequences renders structures in the dataset more distinct from one another and reduces pattern overlaps between folds. For another, the removal of redundant sequences greatly reduces the unbalancedness in the dataset. Some proteins such as myoglobin have been much more extensively studied than others for historical or scientific reasons and consequently there are many more structures for them in the PDB than for others. As a result, the numbers of structures in different folds are vastly unbalanced (see the red curve in Fig 7). This hinders training. At 30% sequence similarity level, the unbalancedness is reduced by about an order (Fig 7).

**Figure 7:**
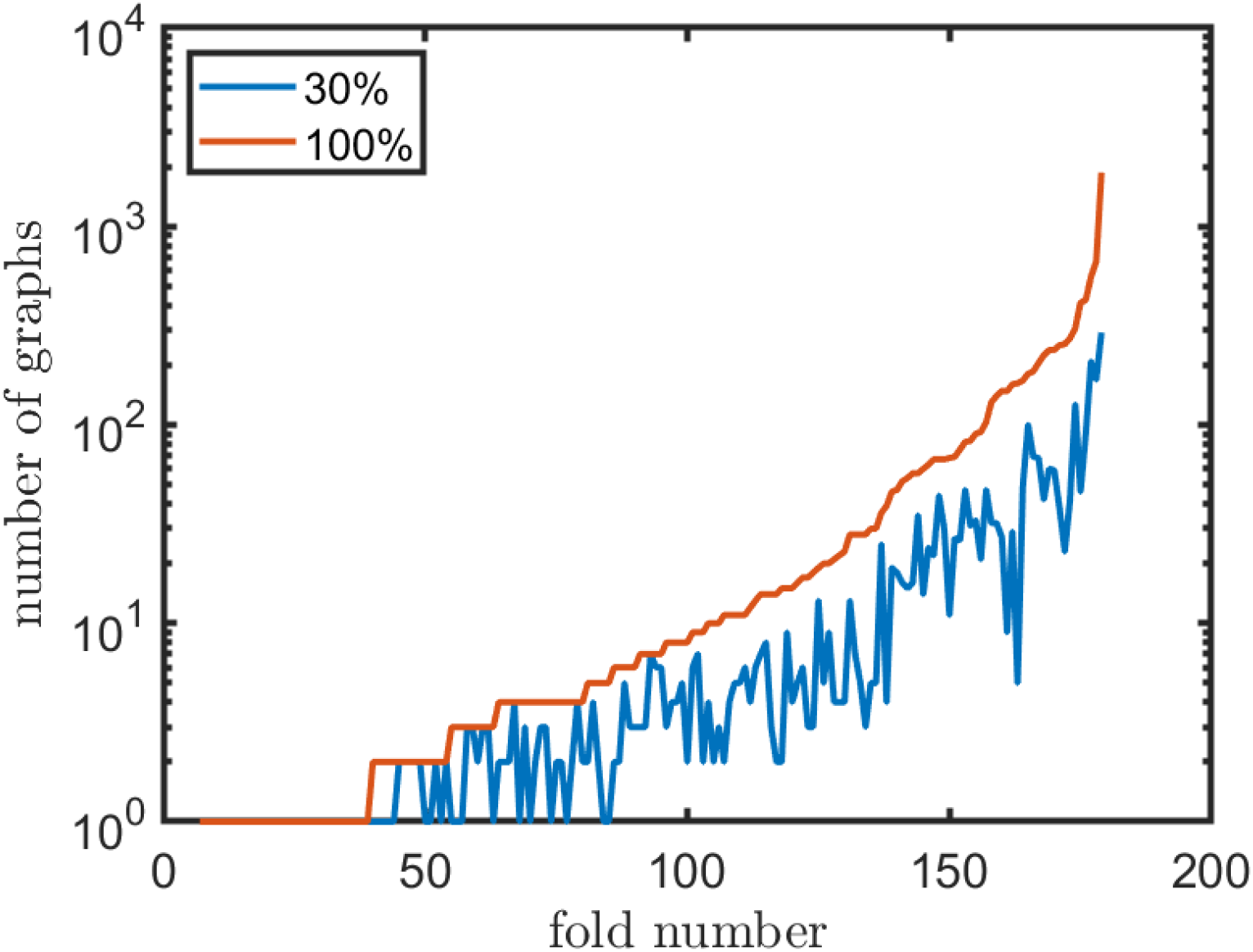
The number of graphs in each fold of the Beta class at different sequence similarity levels: 100% (in red, sorted in the ascending order) and 30% (in blue). The extent of variations in the number of graphs is greatly reduced at 30% sequence similarity (in blue) compared to that at 100% (in red). The graph data with different fold labels are more balanced at a lower sequence similarity level (30%).

The only exception to the above observation is the Alpha class: the accuracy drops to at 82-83% when the the sequence similarity level is reduced to 30-40%. A possible explanation is that compared to the Beta class, the Alpha class has significantly more folds but a similar number of graphs, while when compared to the Alpha/Beta or Alpha+Beta classes, the Alpha class has only one type of secondary structure and thus less diversity in its graphs, which makes it more difficult to train.

### 3.3 Results on CATH Data Set

In addition to the SCOPe 2.07 data set, We have also analyzed the latest release of the CATH data set ^7^. CATH has five levels in their classification hierarchy, namely:

- Class, C-level;
- Architecture, A-level;
- Topology (Fold family), T-level;
- Homologous Superfamily, H-level;
- Sequence families, S-level;

with the counts of Architectures, Topologies, and Super-families in three major classes given in Table 5.

**Table 5:** The counts of Architectures, Topologies, and Superfamilies in three major classes of the CATH dataset.

| Class | No# of Architectures | No# of Topologies | No# of Superfamilies |
| --- | --- | --- | --- |
| 1 (Mainly Alpha) | 5 | 404 | 2033 |
| 2 (Mainly Beta) | 21 | 244 | 1290 |
| 3 (Mainly Alpha Beta) | 14 | 634 | 2337 |

CATH topologies are structurally equivalent to SCOP folds. Assuming that the class and architecture assignments are given/known, we classify structures at the topology level for the CATH dataset. In our first setup, as we did with the SCOPe analysis, we choose not to filter structures by sequence similarity and generate a comprehensive data set that captures the original diversity of the CATH database. The result is given in Table 6. The main difference from the SCOPe analysis is that here we classify topologies under two layers of abstraction, namely Class and Architecture, which makes the classification task comparatively easier. We again apply ten-fold cross validation and train the network for 3, 500 epochs for every dataset under every structural class. The results are given in Table 6. It is seen the prediction accuracy is higher than SCOPe, reaching even 100% for some classes/architectures.

**Table 6:** Prediction accuracies across different topologies of the CATH dataset.

| Class | Architecture | No# of Folds | No# of Domains | Accuracy(%) |
| --- | --- | --- | --- | --- |
| 1 (Mainly Alpha) | 10 (Orthogonal Bundle) | 290 | 53279 | 93% |
| 1 (Mainly Alpha) | 20 (Up-down Bundle) | 104 | 22127 | 95% |
| 1 (Mainly Alpha) | 25 (Alpha Horseshoe) | 6 | 3229 | 98% |
| 2 (Mainly Beta) | 10 (Ribbon) | 26 | 2966 | 98% |
| 2 (Mainly Beta) | 20 (Single Sheet) | 21 | 1335 | 100% |
| 2 (Mainly Beta) | 30 (Roll) | 40 | 7133 | 96% |
| 2 (Mainly Beta) | 40 (Beta Barrel) | 48 | 26846 | 95% |
| 2 (Mainly Beta) | 60 (Sandwich) | 44 | 53682 | 93% |
| 2 (Mainly Beta) | 70 (Distorted Sandwich) | 18 | 4562 | 96% |
| 2 (Mainly Beta) | 105-140 (Beta Propeller) | 6 | 1050 | 98% |
| 3 (Mainly Alpha Beta) | 10 (Roll) | 60 | 15626 | 96% |
| 3 (Mainly Alpha Beta) | 20 (Alpha-Beta Barrel) | 18 | 17355 | 96% |
| 3 (Mainly Alpha Beta) | 30 (2-Layer Sandwich) | 224 | 58283 | 93% |
| 3 (Mainly Alpha Beta) | 40 (3-Layer(aba) Sandwich) | 126 | 90214 | 95% |
| 3 (Mainly Alpha Beta) | 50 (3-Layer(bba) Sandwich) | 11 | 4377 | 100% |
| 3 (Mainly Alpha Beta) | 55 (3-Layer(bab) Sandwich) | 6 | 461 | 100% |
| 3 (Mainly Alpha Beta) | 60 (4-Layer Sandwich) | 16 | 12205 | 97% |

Table 7 presents the results of fold classification of the CATH dataset at 30% sequence similarity level. A comparison between the results from Table 7 and Table 6 reveals again the effect of removing redundant sequences on classification accuracy. At 30% sequence similarity level, the dataset becomes more balanced (Fig 7), which may a the major contributing factor to the higher accuracy seen for most of the architectures, except for Orthogonal Bundle and Up-down Bundle, in Table 7). The significant reduction in the numbers of graphs at 30% sequence similarity level (compared to those at 100% in Table 6) may have further facilitated the convergence of the model during training. Compared to the SCOPe dataset (Table 4), the classification accuracy is relatively higher, probably due to the extra Architecture layer in the CATH classification hierarchy.

**Table 7:**
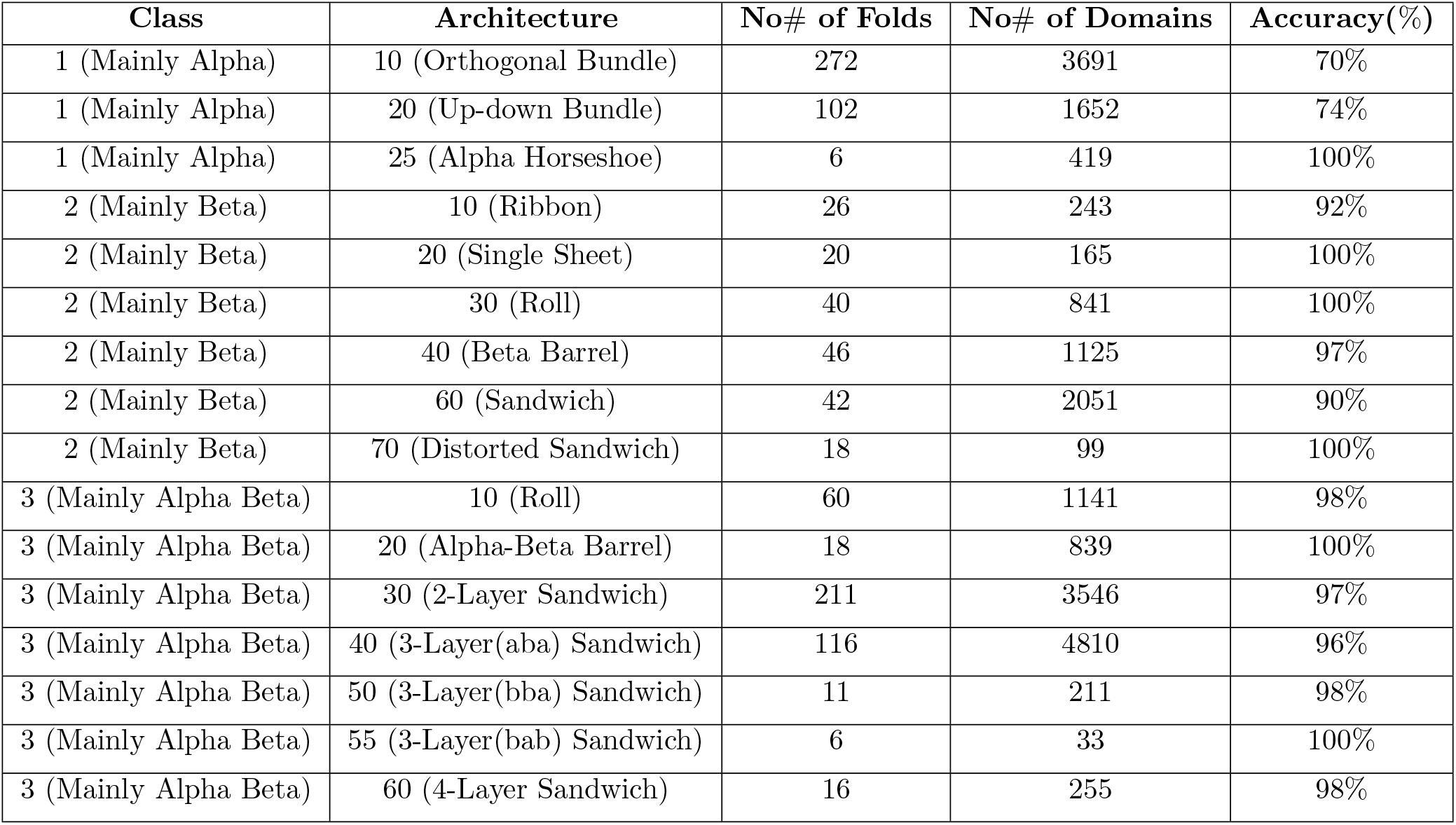
Prediction accuracy across different topologies of the CATH dataset at 30% sequence identity.

### 3.4 Results on Independent Test Data Set

In addition to *k*-fold cross-validation, we performed independent tests on some CATH topologies to assess the classification accuracy of our model on independent test data sets. For this experiment, we drew training samples from three different architectures of three different classes and five topologies under every architecture. A full description of the data set and the results are given in Table 8. We made sure that the training and testing data sets are completely separate and independent and only structures with sequence identity *<*= 30% were selected.

**Table 8:** Results of independent data sets: classification accuracy across three different topologies at 30% sequence identity.

| Class | 1 (Mainly Alpha) | 2 (Mainly Beta) | 3 (Mainly Alpha Beta) |
| --- | --- | --- | --- |
| Architecture | 10 (Orthogonal Bundle) | 70 (Distorted Sandwich) | 40 (3 layer(aba) Sandwich) |
| Topologies | 8,10,12,20,30 | 98,100,160,170,240 | 5,20,30,47,50 |
| No# of Training | 353 | 335 | 416 |
| No# of Testing | 56 | 45 | 65 |
| No# misclassification | 8 | 6 | 4 |
| Accuracy(%) | 48/56 = 86% | 39/45 = 87% | 61/65 = 93% |

This is a multi-class classification problem and we have done all-vs-all classification here. In this analysis, we computed the accuracy of the classifier using the standard *Q* percentage accuracy ^28^ with *Q* = *C/N*, where *Q* is the total accuracy, *C* is number of correct classification, and *N* is total number of test samples combining all the classes.

### 3.5 Comparison with Structure Alignment Methods

In this section, we compare our graph neural network model for fold classification with the state-of-the-art structure alignment approaches such as TMalign ^29^ and mTMalign ^30^. mTMalign produces multiple structure alignment based on TM-score of paiwise structure alignments. Both mTMalign and TMalign rely on pairwise sequence similarity between structures. A recent study by Holm ^31^ using 140 query domain structures and the SCOPe dataset showed that structure alignment-based approaches are rather poor for fold detection when structures are distantly related, i.e., when the database contains no structures of the same family or super-family as the query domain structure.

Since the focus of the present work is fold classification, which is slightly different from fold detection, we use Holm’s results in the following way for fold prediction/assignment. For each query domain, we examine the top 10 structural matching predictions from mTMalign/TMalign ^30 29^, excluding matches that are from the same superfamiliy as the query domain structure. Based on the top 10 matches and their corresponding SCOPe fold assignments, We a perform majority voting to determine the fold assignment for the query domain. Doing this allows us to assess how well structural matching approaches can predict folds when only distant superfamilies are available (for the query protein). We then compare the results with the fold predictions by our graph neural network based approach. To have a fair comparison between our model and structure-matching methods, we exclude from our training set all structures that belong the same superfamilies as the query domains. We then test our trained model with the query domains to check the number of true positives. For example, query domain d1cuka1 from all alpha class belongs to long alpha-hairpin fold. This fold spreads into 20 superfamilies. In our training set, we exclude the superfamily of d1cuka1 and include only the remaining 19 superfamilies in the training.

The results, in terms of the number of true positives, are given in Table 9. For example, for mTMalign, there are 4 true positive cases out of the 11 for the Alpha class, and 2 out of 14, 4 out of 12, and 0 out of 5 for the remaining three classes, respectively.

**Table 9:** A comparison between our ML-based approach and two structure matching-based approaches (mTMalign and TMalign) in fold prediction for query structural domains that belongs to a new superfamily. The number of query domain structures used in each class is listed in the top row. The table lists the accuracy in fold assignments for the query domains.

| Class (# of queries) | Alpha (11) | Beta (14) | Alpha+Beta (12) | Alpha/Beta (5) |
| --- | --- | --- | --- | --- |
| **our approach** | 5/11 | 4/14 | 3/12 | 2/5 |
| **mTMalign** | 4/11 | 2/14 | 4/12 | 0/5 |
| **TMalign** | 2/11 | 2/14 | 3/12 | 1/5 |

Next we repeat the same analysis but keep every structure in the dataset except the query domain structures. This corresponds to an easier fold prediction scenario where for the query domain, there exist in the dataset domain structures of the same family/superfamily. It was known that for such a scenario structure matching approaches perform extremely well in fold detection ^31^. Indeed, this is confirmed in the fold assignment accuracy results given in Table 10.

**Table 10:**
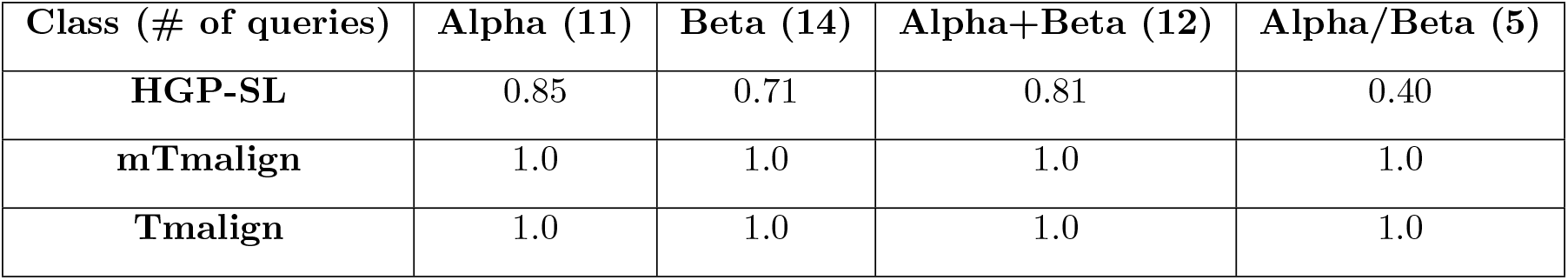
A comparison between our ML-based approach and two structure matching-based approaches (mTMalign and TMalign) in fold prediction for query structural domains that belong to an existing family or superfamily. The number of query domain structures used in each class is listed in the top row. The table lists the accuracy in fold assignments for the query domains.

The results show that for structure matching methods, the accuracy in fold assignment for a domain that belongs to an existing family or superfamily increases dramatially. As we consider the top 10 hits for every query domain, we see that structures of the same family or superfamily are nearly always selected by structure matching methods (namely TMalign and mTMalign) and thus produces an accuracy of 100%. For our neural network-based model, the improvement in accuracy is not as dramatic. It is possible, however, when we include more node level features, further improvement in accuracy could be achieved. One important observation here is that the inclusion of structures of the same family/superfamily certainly improves the training and fold assignments. Compared with structure matching based methods, our deep neural network model is less dependent on the existence of structures of the same family or superfamily. This makes our approach more suited for fold assignments for query domains of a novel superfamily. Our approach may also be adapted to identify a new fold.

### 3.6 Summary

There are a few interesting observations after analyzing both CATH and SCOPe 2.07 data sets.

- For both CATH and SCOPe data set, we can see that the all alpha class got lower accuracy compared to others. At different sequence similarity levels their accuracy ranges from 88% − 96% compared to other classes like beta and alpha/beta. One reason could be that alpha fold patterns are less distinct from one another, which is reflected on their overall graph representations. This makes their classification more challenging than other classes.
- The SCOPe data set is most difficult when the data set is large and there are many different folds in the data set. SCOPe groups structures into a few structural classes and diverse superfamilies may be placed under the same fold. This makes it harder to train and optimize, especially when the data set is large.
- For the CATH data set, there is an additional level in the hierarchy between the class and topology levels, namely the architecture level. Under one architecture, structures may share a common overall shape but different topologies. Although CATH topology is the counterpart of the SCOPe fold, the topologies in CATH are more finely categorized by their architectures, making it easier to learn patterns for generalization from CATH. This is seen from the results in Table 7: even when a certain architecture/class contains a large number of topologies and consequently a large graph data set, the classification accuracy remains high. Whereas for SCOPe, the classification accuracy is always affected by the size of the data set - the more graphs are in the data set, the lower is the accuracy.

### 3.7 Misclassification Analyses

Results on the CATH and SCOPe data set show that certain topologies get a lower accuracy than others. To analyze the patterns of misclassification, we examine closely some misclassification cases and compared them with their actual classification by visualizing both their structures and their topology graphs. We visually examine these structures in PyMol to see if there is any obvious reason for the misclassification. Figure 8 shows the protein graphs of two structures from the mainly-alpha-beta roll architecture. PDB entries are 1yrw chain A (or 1yrwA) of CATH topology 25 (Figure 9(A)), and 2he7 chain A (or 2he7A) of CATH topology 20 (Figure 9(B)). In this case, 1yrwA is misclassified as topology 20. We explored all the alpha beta graphs from topology 20 and found out that PDB entry 2he7A of topology 20 has a similar topological arrangement like 1yrwA of topology 25 (see Figure 9). Both entries have ten alpha helices and 11 and 12 beta strands. This similarity explains why 1yrwA is misclassified into the topology of 2he7A.

**Figure 8:**
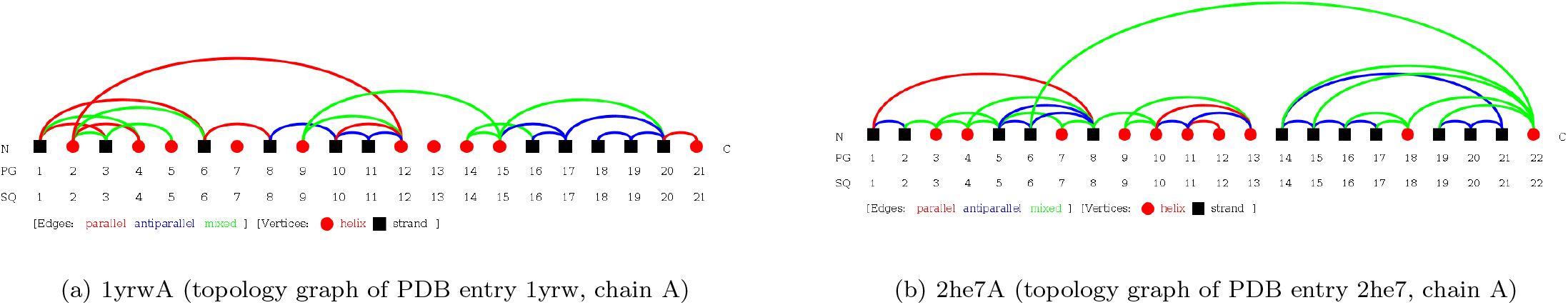
Topology graphs of two alpha-beta-roll proteins

**Figure 9:**
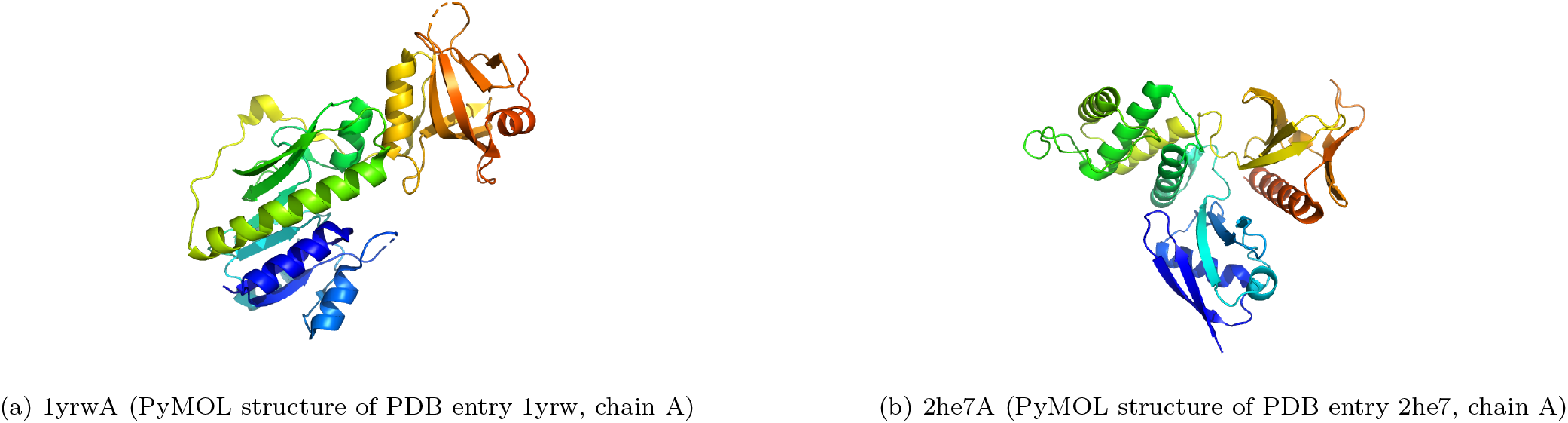
PyMOL structures of two alpha-beta-roll proteins

We further explore another classification error from the orthogonal-bundle architecture of the mainly-alpha class. The PDB entry 1mu5 chain A contains a domain belonging to CATH topology 8 of the orthogonal bundle architecture. This domain is misclassified as topology 135. When compared with the other PDB entries from topology 135 such as 3ju5 chain A, we find that 3ju5A is highly similar to 1mu5A in terms of the number of helices in their alpha graphs (Figure 10). Both entries have 18 helices and a similar overall secondary structure composition (Figure 11), which explains the misclassification.

**Figure 10:**
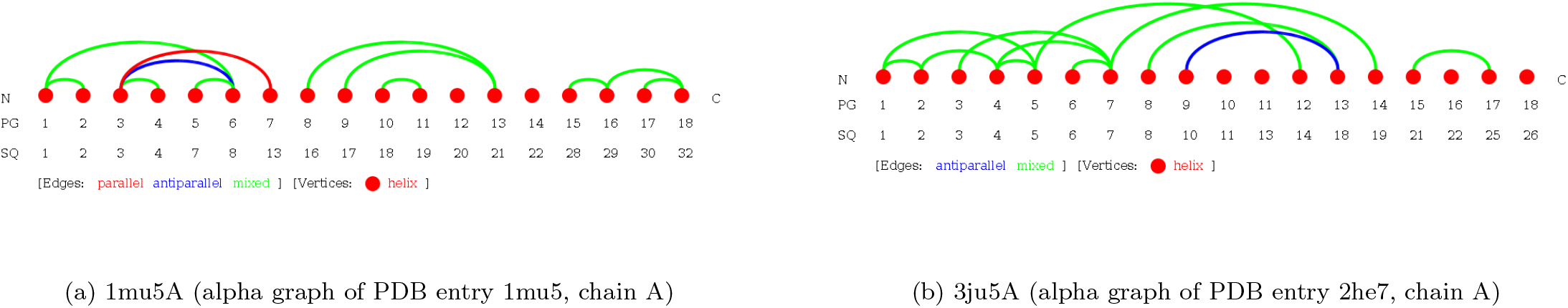
Topology graphs of two mainly-alpha-orthogonal-bundle proteins

**Figure 11:**
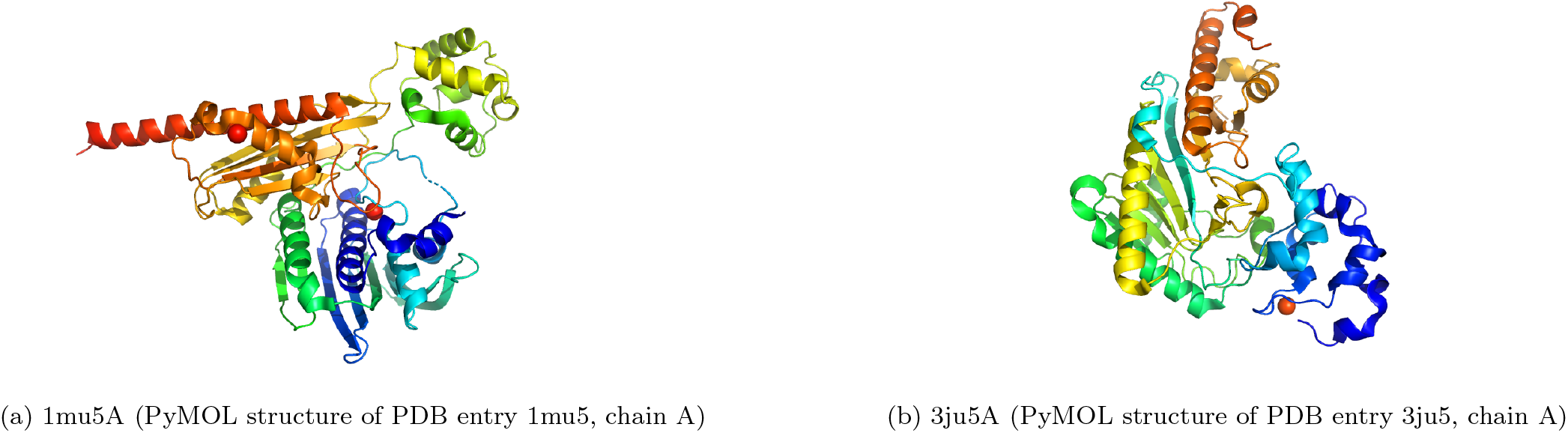
PyMOL structures of two mainly-alpha-orthogonal-bundle proteins

## 4 Materials and Methods

### 4.1 Protein Structure Classification

Every protein structure has an evolutionary origin and can be classified into their taxonomic hierarchy. Also to better understand the structural relationship with other proteins, their taxonomic hierarchical classification gives important information. SCOPe and CATH are two databases that classify protein structure topologies into their taxonomic hierarchy. Although these traditional approaches are present, automated approaches are in need due to the rapid growth in the availability of new structures. Specifically, machine learning-based approaches for structure classification are becoming increasingly attractive. SCOPe is a mainly manually curated database that classifies protein structures in their fold hierarchy considering their evolutionary relationship. At first, SCOPe partitions proteins into domains These domains are then classified into four levels. The highest level of this hierarchy is class. There are seven classes in SCOPe, namely, (a) alpha proteins, (b) beta proteins, (c) alpha/beta proteins, (d) alpha+beta proteins, (e) multi-domain proteins, (f) membrane and cell surface proteins and peptides, and (g) small proteins. Class describes the content of secondary structure elements in protein domains. The next level in this classification hierarchy corresponds to protein fold. Fold groups structures having similar number and order of secondary structure elements with similar connectivity patterns. There are 1232 protein folds all together in SCOPe. Fold groups together structurally similar superfamilies that might not have any evolutionary relationship. Family contains proteins that share similar sequences and close evolutionary relationships. Superfamily combines proteins with similar functional and structural features that descend from same ancestor. The primary concern of this work is to classify protein structures into their respective folds. In summary it can be said that, the evolutionary relationship among known protein structures is the basis of classification for SCOPe. On the other side, CATH classification relies on structural rearrangements and structural properties of proteins. It applies a structure comparison-based method for classifying unknown structures. CATH has three major classes in their first layer. Those are: a) mainly alpha, b) mainly beta and c) mainly alpha-beta. In the second layer, CATH classifies the structures based on the common overall shape of the topologies. This level is called the architecture level. Combining the three major classes there are around 40 architectures in CATH. The next level is the topology/fold level. Inside topology level, structures are clustered based on their architecture and connectivity pattern of the secondary structure elements. Combining the 3 major classes and 40 architectures, there are around 1282 topologies in CATH. One interesting observation here is, although CATH classifies much more domains than SCOPe, the numbers of fold/topology in both databases are almost equal. It reminds that the number of folds in nature is limited although the structure and sequences are ever increasing. Finally, after the topology layer, CATH classifies the structures into homologous superfamiles. There are around 5660 superfamilies in the CATH database, combining all the structural classes, architectures and folds. CATH superfamily is a mixture of SCOPe superfamily and family and the number of superfamilies in CATH reflects the sum of both.

### 4.2 Protein Topology Graph Library

The PTGL comprises protein topologies based on unique graph-theoretic definitions at different levels of abstraction. Protein topology graphs were defined on the secondary structure level mainly for protein structure prediction ^32^, analysis ^33^ and comparison of protein structures ^34,35^.

PTGL ^19 16 36^ reads the atom coordinates from a macromolecular crystallographic information file (mmCIF) of the PDB ^37^ and the assignments of amino acids to secondary structure elements from a DSSP file ^38^, all per chain. All types of helices are treated as one helix type. Single beta bridges that occur next to a beta sheet are treated as beta sheet. Secondary structure elements that are interrupted by one differing classification, e.g. loop, are merged. Finally, PTGL neglects secondary structure elements of less than three amino acids.

PTGL computes all atom-atom contacts applying a Euclidean distance less than 4 Å. Depending on the involved atoms, the contacts are classified as backbone-backbone (BB), backbone-sidechain (BC) or sidechain-sidechain (CC). A contact between secondary structure elements is defined in dependence of the type of involved secondary structure elements, i.e., helix or strand, and rules for the involved atom contacts, i.e., BB, BC or CC (see Table 11).

**Table 11:** Required residue-residue contacts for assignment of a contact between secondary structure elements. Residue-residue contacts are differentiated by the type of the atoms involved in the contact as backbone-backbone (BB), backbone-sidechain (BC) and sidechain-sidechain (CC) contacts.

| Type of SSE 1 | Type of SSE 2 | Required residue-residue contacts |
| --- | --- | --- |
| beta strand | beta strand | BB > 1 or BC > 2 |
| beta strand | helix | BB > 1 or BC > 3 or CC > 3 |
| helix | helix | BC > 3 or CC > 3 |

A direction of secondary structure elements can be inferred from N to C terminus. PTGL creates a vector in three-dimensional space for each SSE. The initial point and terminal point are the geometric mean of the atom positions of the first and last four residues, respectively. For shorter secondary structure elements of length *n*, the first and last *n* − 1 residues are used instead. PTGL computes for each secondary structure element contact the angle between the secondary structure elements and assigns a parallel, mixed or antiparallel orientation of the secondary structure elements to each other for angles up to a degree of 65, 115 and 180, respectively.

We define a Protein Graph (PG) of a protein chain, where the vertices correspond to secondary structure elements and the edges to spatial neighborhoods of the vertices (see Figure 12a). The vertices are labeled by their SSE type, i.e., strand or helix. The edges are labeled by the orientation of the participating secondary structure elements to each other, i.e., parallel, mixed or antiparallel. A Folding Graph (FG) is a connected component of a PG (see Fig. 12b, vertices 2 - 4). An alpha, beta or alpha-beta (albe) PG or FG contains only helices, only strands or both helices and strands, respectively (see Fig. 12).

**Figure 12:**
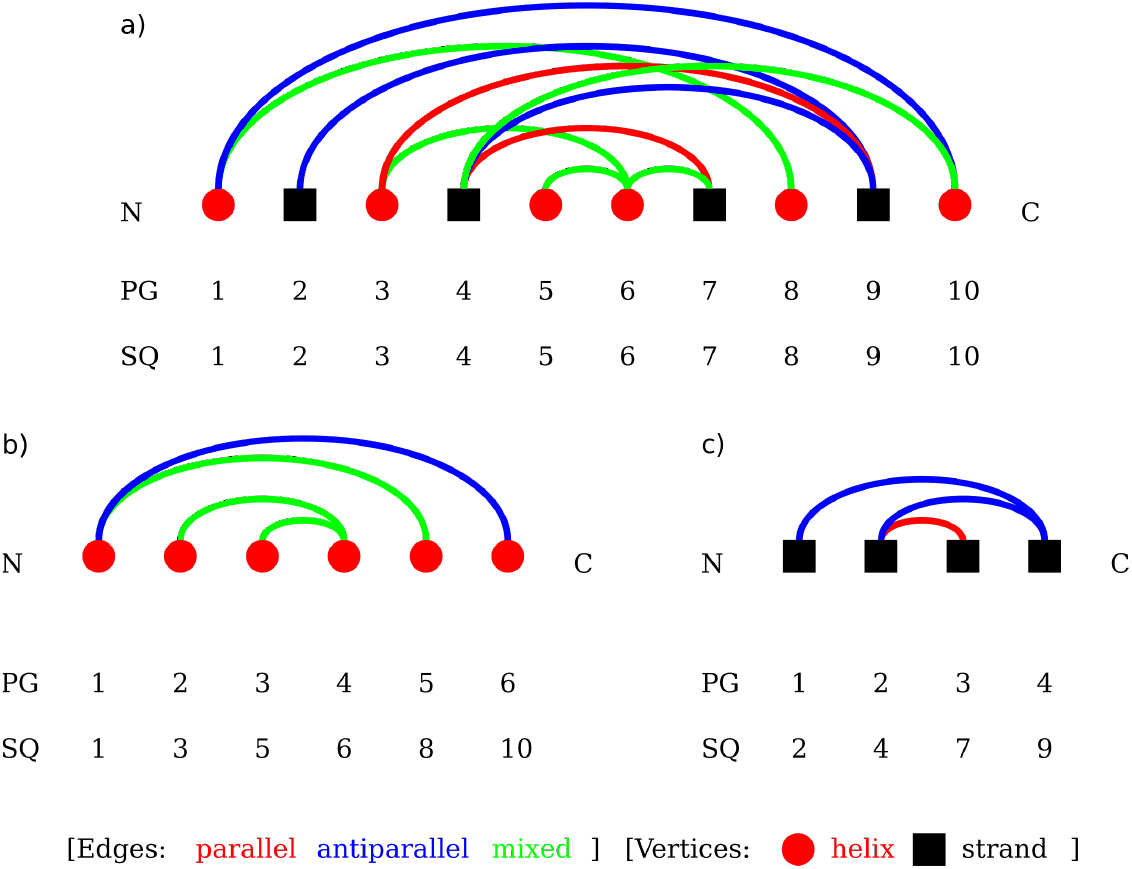
A (a) alpha-beta, (b) alpha and (c) beta Protein Graph. The vertices are horizontally aligned from N to C terminus. Helices and strands are drawn as red circles and black boxes, respectively. Edges represent spatial neighborhoods and are colored red, blue or green for parallel, antiparallel and mixed orientation of the involved secondary structure elements to each other, respectively. The vertices are labeled with increasing IDs by occurrence in the graph (PG) and occurrence in the sequence (SQ).

**Figure 13:**
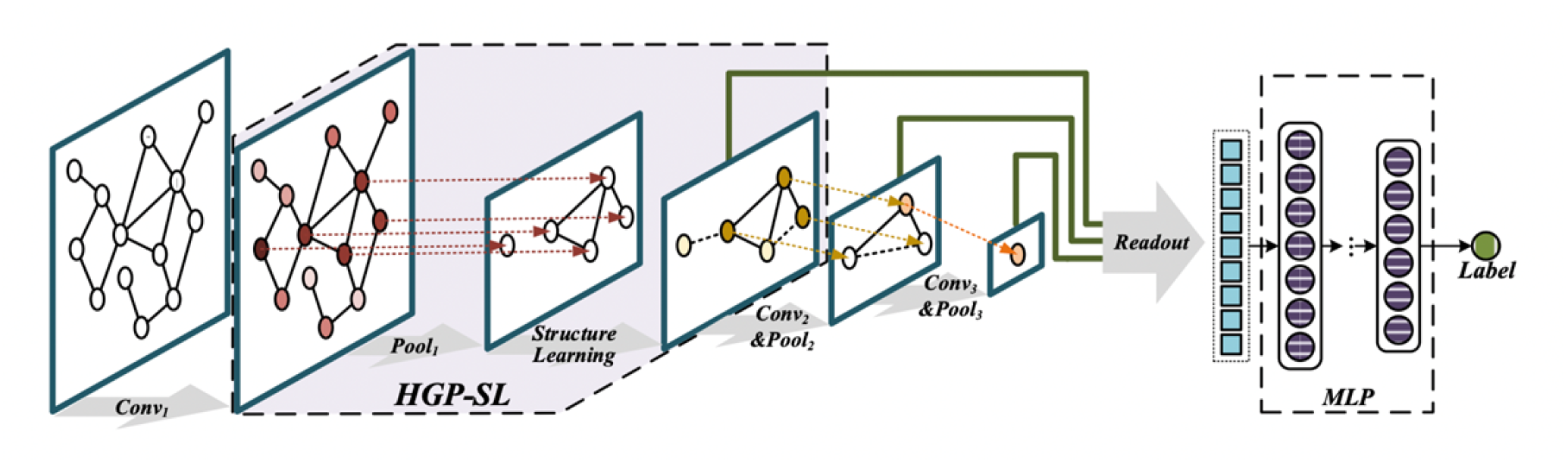
Hierarchical Graph pooling with Structure Learning for Graph Classification. https://github.com/cszhangzhen/HGP-SL

### 4.3 Graph Classification

Graph classification is the problem of learning a function from a set of attributed graphs to predict their class labels. Given a set of attributed graphs *G* = {(*G*1, *y*_1_), (*G*_2_, *y*_2_), …, (*G*_*n*_, *y*_*n*_)}, the goal is to learn a function *f: G* → *Y* to approximate the graph labels. Here, *G* is the input space of the graphs, and *Y* is the set of all graph labels.

### 4.4 Graph Processing and Feature Extraction

We identified the PTGL graph in JSON format corresponding to every entry in every data set and then parsed that graph to convert into a pyTorch geometric graph. The JSON-formatted graph contains all the relevant information regarding the graph structure, such as node labels, edge labels, adjacency relations etc. We parsed and processed these information to capture the node features and construct the graph in pyTorch format from the adjacency relation. In this work, we considered only node features for every pyTorch graph. We have selected three node features to learn patterns from the graph. Those are the node types; (alpha helix or beta strand), node degrees and the number of residues or length of secondary structure element element in each node.

### 4.5 HGP-SL for Protein Fold Classification

The HGP-SL (Hierarchical Graph Pooling with Structure Learning) work was developed by Zhang et al. ^15^. A summary of HGP-SL is given below. Readers are referred to the original work ^15^ for details.

#### 4.5.1 Problem Formulation

Given a set of labelled graph data *G* = {*G*_1_, *G*_2_, …, *G*_*n*_}, the goal is to learn patterns from this data set and predict the label for unknown graphs. Each graph could be represented as *G*_*i*_ = (*V*_*i*_, *E*_*i*_, *A*_*i*_, *X*_*i*_, *Z*_*i*_, *Y*_*i*_). *V*_*i*_ and *E*_*i*_ represent the node and edge set of the graph, where *n*_*i*_ and *e*_*i*_ correspond to the number of nodes and edges, respectively, in the graph. 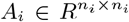 represents the adjacency matrix which contains all the edge information of the graph. 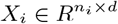 is the node feature matrix where *d* represents the dimension of the node attributes. 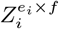 is the edge features matrix where *f* is the dimension of the edge attributes. *Y* ∈ *R*^*n×s*^ is the label matrix that corresponds to the full graph data set. Here, *s* is the length of the label set, and *n* is the number of graphs in the data set. *Y*_*i*_ contains the label for an individual graph *i*. During the graph pooling operation, the number of nodes and graph structure changes at each subsequent layer. The *i*th graph trained on *k*th layer could be represented as 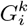 having nodes 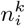. The adjacency matrix and hidden representation matrix for the *i*th graph at *k*th layer could be represented as 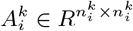 and 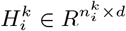.

#### 4.5.2 Graph Convolutional Neural Network

Graph Convolutional Neural Networks (GCN) are a variant of GNN. The graph convolution operation can be categorized into two branches. Those are spatial and spectral approaches. The spectral-based approaches define graph convolution operation by parameterized filters from the perspective of graph spectral theory ^39^. However, due to the computational flexibility and efficiency, spatial-based approaches gained much popularity in recent years. Spatial-based approaches are analogues to conventional convolution operation on images. Graph convolution works by utilizing node’s spatial relation. By directly aggregating node’s neighborhood information, node-level representations are generated in every subsequent convolution layer, which in turn contributes to generate the graph-level representation. For generating a representation for *k*th layer, GCN takes as input graph *G*’s adjacency matrix and *k* − 1th layer’s hidden representation matrix. It generates the representation for *k*th layer as follows:

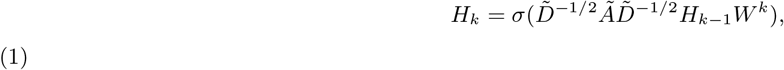

where *σ* is the non-linear activation function. *Ã = A + I* is the adjacency matrix with self-connections. 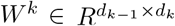 is the trainable weight matrix for layer *k*, and 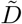 is the diagonal degree matrix of *Ã*.

#### 4.5.3 HGP-SL

HGP-SL ^15^ is a novel graph classification approach with sophisticated pooling operation. This method utilizes the node features and the entire graph’s topological information for predicting labels associated with the graphs. The common approach for graph classification is to summarize all node information to create a global representation of the graph. This approach neglects the entire graph’s structural information, and therefore, the graph level representation that is generated this way is not very intense. On the other hand, GNN cannot aggregate node information in a hierarchical way. Moreover, to create a meaningful representation of the graph, sometimes it is required to put a special significance to graph substructures, as different substructure may contribute differently to the whole graph representation. That’s why, to learn from both graph local information and global structure and to generate hierarchical representation, a graph pooling operation is required. HGP-SL provides such a pooling process to meet this requirement. It provides a sophisticated pooling layer in addition with a graph convolution layer to be used with different graph neural network architectures. In this subsection, we will describe the two major components of HGP-SL for graph classification.

- **Graph Pooling Operation:** The pooling layer in HGP-SL is a non-parametric layer. The proposed pooling layer works by selecting a set of informative nodes to produce an induced sub-graph for the next layer. The selection process works as follows:
  - It first re-orders the nodes according to the node information scores.
  - The node information score can be computed by the Manhattan distance between the node representation itself and the one calculated from its neighbors:

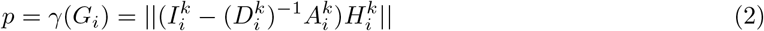 Here, 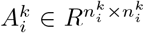 and 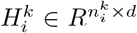 are the adjacency and node representation matrices, respectively. 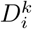 is the diagonal matrix of 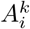, and 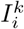 is the identity matrix. 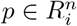 encodes the information score of each node in the graph. *γ* is the node score-encoding function. Here, it is the Manhattan distance.
  - After calculating the node information score of every node, top-ranked nodes are selected as follows:

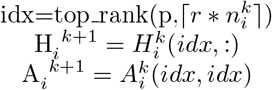 Here *r* is the pooling ratio, *top_rank* is the function by which top 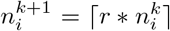 indices are selected. For the induced sub-graph, the node representation matrix and adjacency matrix are extracted using 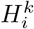(*idx*, :) and 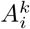(*idx, idx*), respectively. So, both the node feature and graph structure information are retained this way for the next layer.
- **Structure Learning Mechanism:** HGP-SL incorporates a novel structure learning mechanism to learn a refined structure from the induced sub-graph in each pooling layer. After the pooling operation, the induced sub-graph structure may result in highly correlated nodes being disconnected, which in turn may prevent message passing and effective node level representation learning. Moreover, the induced sub-graph structure might preserve important domain knowledge for the graph that originates from a specific domain-like protein structure. Considering all these issues, the structure-learning mechanism turns out to be an effective scheme to produce meaningful hierarchical representation of the graph. The basics of the structure-learning layer is a single-layer neural network parametrized by a weight vector 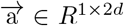. It takes as input graph *G*_*i*_’s *k*th hidden layer presentation 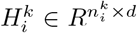 and structure information 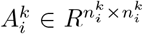 and encodes a refined graph structure by learning the pairwise relationship between each pair of nodes using an attention mechanism. The similarity score between two nodes *v*_*p*_ and *v*_*q*_ can be expressed by the following equation:

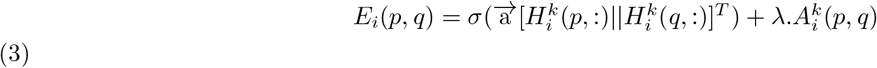

Here, *σ* is a non-linear activation function such as ReLU. 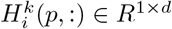 and 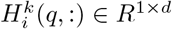 correspond to the *p*th and *q*th row of the hidden representation matrix 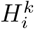 that represent the nodes *v*_*p*_ and *v*_*q*_, respectively.

#### 4.5.4 Protein Fold Classification

We formulate the protein fold classification problem as a graph classification problem according to the definition described in section 4.3. Here, we take as input, the protein graphs of PTGL for a number of PDB entries from SCOPe and CATH data sets and give them labels according to the SCOPe/CATH fold labels.

### 4.6 Data Sets

We have analyzed the SCOPe 2.07 data set for training, validation and testing. We classify four different structural classes of proteins, namely, all-alpha, all beta, all alpha and beta proteins (a/b), all alpha and beta proteins (a+b). We also examine the latest version of the CATH data set. CATH classifies the proteins in three major classes; mainly-alpha, mainly-beta and mainly-alpha-beta. Additionally, we examine three different fold recognition benchmark data sets, namely the DD data set ^27^, EDD data set ^21^, and TG data set ^40^. The DD data set consists of a training and testing data set. The sequences in this data set cover 27 protein folds, which are categorized into four protein structural classes; alpha, beta, alpha/beta, alpha+beta. The training set contains 311 protein sequences with ≤ 40% residue identity, and the testing set contains 383 sequences with ≤ 35% residue identity. Also the training and testing set have ≤ 35% sequence similarity. The EDD data set is an extended version of DD data set, which is also composed of 27 protein folds categorized into four different classes. There are 1612 protein sequences in the TG data set, which covers 30 different folds from SCOP 1.73 having ≤ 25% sequence similarity. These three benchmark data sets are classic data sets made up with sequence data, for which we constructed topology graphs for our graph classifier and compared the results with the sequence-based models.

#### 4.6.1 Network Parameters

We trained the HGP-SL model on all the aforementioned benchmark data sets in addition to the SCOPe 2.07 data set. We tried different network settings for different data sets. A batch size of 512 and number of epoch of 3000 was mainly used. The learning rate, dropout ratio and pooling ratio were 0.005, 0.1 and 0.5 most of the time. During the training time, the model optimizes by generating minibatches of graphs, where each batch is considered as a connected graph and within this densely connected graph, each individual graph is like a connected sub-graph.

### 4.7 Evaluation Strategy

We applied ten-fold cross validation in our data set. We divided each data set into ten equal-sized groups and used eight out of ten for training, one for test and one for validation. We did this for every unique group. That means, every data sample got the opportunity to be used in the holdout or test set once. The model is trained ten times, and then, the overall score is computed by taking the mean of the ten scores.

## 5 Conclusion

In this study, domain specific protein graphs from PTGL and graph neural network (GNN) have been used for protein fold classification. The work demonstrates that graph neural network is potent for fold assignments, especially for query proteins of a novel superfamily, in which regard it outperforms even top-notch structure-matching-based methods. As a future work, we plan to extend our method to classify protein complexes.

